# Complete correction of murine phenylketonuria by selection-enhanced hepatocyte transplantation

**DOI:** 10.1101/2023.08.27.554228

**Authors:** Anne Vonada, Leslie Wakefield, Michael Martinez, Cary O. Harding, Markus Grompe, Amita Tiyaboonchai

## Abstract

Hepatocyte transplantation for genetic liver diseases has several potential advantages over gene therapy. However, low efficiency of cell engraftment has limited its clinical implementation. This problem could be overcome by selectively expanding transplanted donor cells until they replace enough of the liver mass to achieve therapeutic benefit. We previously described a gene therapy method to selectively expand hepatocytes deficient in cytochrome p450 reductase (Cypor) using acetaminophen (APAP). Because Cypor is required for the transformation of APAP to a hepatotoxic metabolite, Cypor deficient cells are protected from toxicity and are able to expand following APAP-induced liver injury. Here, we apply this selection system to correct a mouse model of phenylketonuria (PKU) by cell transplantation. Hepatocytes from a wildtype donor animal were edited in vitro to create Cypor deficiency and then transplanted into PKU animals. Following selection with APAP, blood phenylalanine concentrations were fully normalized and remained stable following APAP withdrawal. Cypor-deficient hepatocytes expanded from <1% to ∼14% in corrected animals, and they showed no abnormalities in blood chemistries, liver histology, or drug metabolism. We conclude that APAP-mediated selection of transplanted hepatocytes is a potential therapeutic for PKU with long-term efficacy and a favorable safety profile.

## Introduction

Hepatocyte transplantation is a strategy that represents a potential therapeutic approach for any genetic disorder affecting the liver(1). Hepatocytes are delivered into the portal circulation, usually by catheterization of the portal or umbilical veins(1). Hepatocytes then engraft in the liver and support liver function long-term. Hepatocyte transplantation has been applied clinically in over 100 human patients with a variety of inborn and acquired liver disorders, in both adult and pediatric patients(2, 3). Although partial phenotypic correction has been described in numerous disorders(1, 4, 5), complete and sustained correction of inborn errors of metabolism has not yet been reported. This lack of complete efficacy is likely due to the low efficiency of engraftment of the delivered cells. Only approximately 1% of the recipient’s liver mass can be replaced through standard transplantation methodology, which is insufficient for complete phenotypic correction for the majority of disorders(6). This limitation could be overcome by providing a selective advantage to allow cell division of the transplanted donor cells until they make up a sufficient percentage of the hepatic mass to allow disease correction. This threshold is estimated to be about 10% for several genetic liver conditions(1).

We have previously reported a robust system for the selective expansion of hepatocytes that were genetically modified in vivo using gene therapy methods(7). This system relies upon the use of the common fever and pain medication acetaminophen (APAP) as a selection agent. APAP is metabolized in hepatocytes through several metabolic pathways. Although a majority is directly converted into non-toxic metabolites, a minority is acted upon by cytochrome p450 enzymes (Cyps) in hepatocytes in zone 3 of the hepatic lobule to form an electrophilic intermediate, *N*-acetyl-*p*-benzoquinone imine (NAPQI)(8). NAPQI is hepatotoxic at high concentrations, depleting hepatocytes of glutathione. The Cyps involved in this process require a cofactor, cytochrome p450 reductase (Cypor)(9). Thus, Cypor knockout prevents Cyp-mediated metabolism and renders targeted cells resistant to APAP-induced hepatotoxicity. We showed in prior work that hepatocytes made Cypor deficient by CRISPR gene editing can be expanded over 100-fold to > 40% of the hepatic mass in vivo by APAP treatment(7, 10).

Here, we apply the APAP selection approach to the selective expansion of transplanted hepatocytes in mouse models of phenylketonuria (PKU). PKU is caused by a deficiency in phenylalanine hydroxylase (Pah). Pah is normally expressed in hepatocytes, and it functions to convert dietary phenylalanine (Phe) into tyrosine. In the absence of Pah, high circulating blood Phe levels lead to a severe neurological phenotype. PKU patients must maintain a severely protein-restricted diet throughout their lifetime to avoid intellectual disability(11). If blood Phe levels are maintained at ≤360 µM by dietary restriction or alternate therapy, neurologic symptoms are largely avoided(11). Several human hepatocyte transplantation trials have shown partial correction of PKU, but complete correction has not yet been achieved(12, 13). Studies in mouse models have demonstrated that approximately 10% replacement of the liver with donor hepatocytes is sufficient for correction of blood Phe levels(14, 15). Here, we demonstrate that APAP selection of transplanted hepatocytes can achieve the therapeutic cell replacement threshold for PKU and provide complete and long-term correction of hyperphenylalaninemia.

## Results

### In vivo selection of mouse hepatocytes for PKU

To prepare donor cells, primary mouse hepatocytes were isolated by collagenase perfusion of the liver from a healthy adult donor mouse(10). The donor animal was wildtype for *Pah*, hence transplanted hepatocytes were capable of Phe metabolism. They also expressed a membranous *tdTomato* marker transgene to allow for in vivo visualization of transplanted cells(16). Hepatocytes were treated in vitro with ribonucleoproteins containing SpCas9 protein and a *Cypor*-targeting single guide RNA (sgRNA) delivered by Lipofectamine. This procedure yielded insertion/deletion mutations (indels) in 84-90% of alleles in the total population of transfected cells (Figure S1), as assessed by analysis of genomic DNA (gDNA) from hepatocytes that were plated after transfection and analyzed using the TIDE algorithm(17).

Hepatocytes were then delivered to adult hyperphenylalaninemic male and female mice homozygous for the *Pah^enu2^* mutation via intrasplenic injection(18). This mouse model has a missense point mutation in the *Pah* gene that results in the production of a non-functional protein(19). Mice were assessed for baseline blood Phe levels 2-4 weeks after transplantation and continued to have elevated blood Phe levels ≥1500 µM, indicating that the transplanted cell dose was subtherapeutic (Figure 1A). Mice were then placed on a selection diet containing 1.5% APAP w/w. Mice were maintained on this diet for 50 days, at which time mice were placed on standard chow for 10 days prior to being assessed for blood Phe concentration. APAP was discontinued in animals that showed corrected blood Phe (≤360 µM) at this time. APAP-treated animals that still showed elevated blood Phe levels after 50 days of selection were placed on APAP diet for an additional 25 days before being analyzed again. At this timepoint, APAP diet was discontinued because biochemical correction had been achieved in all animals. Following APAP treatment, blood Phe concentrations decreased to within the corrected range in both male and female mice to an average of 207 µM ± 78 µM and as low as 130 µM. Blood Phe levels remained within the corrected range for more than 250 days after APAP diet was stopped (Figure 1A). Control animals that received cell transplantation but remained on standard diet without APAP showed highly elevated Phe levels throughout the course of the experiment. Corrected mice of both sexes also showed the characteristic darkening of coat color associated with a decrease in blood Phe (Figure 1B, Figure S2A). A significant increase in body weight was also seen in corrected male mice as compared to unselected controls (Figure S2B).

**Figure 1:**
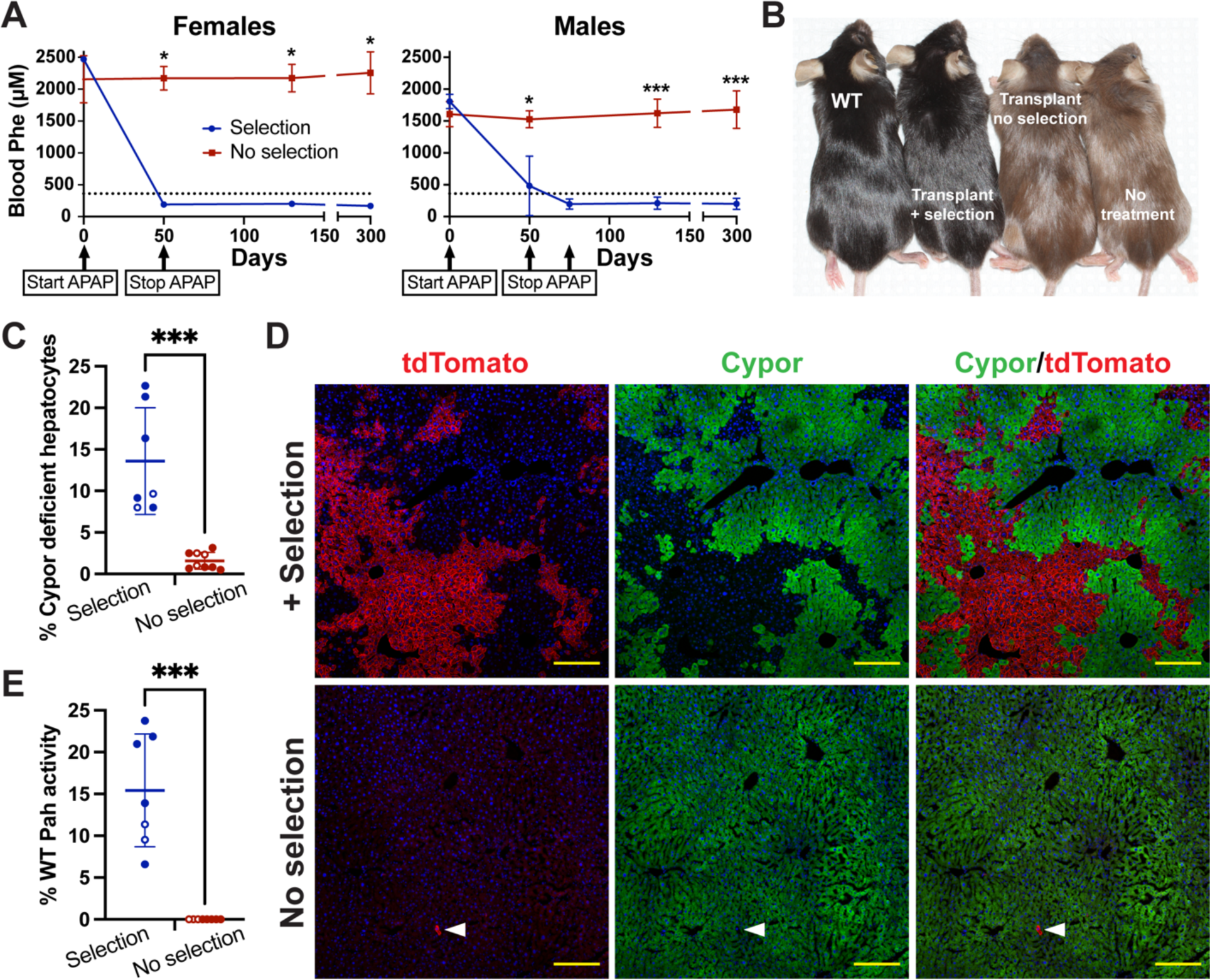
APAP selection of transplanted hepatocytes in *Pah^enu2/enu2^* mice. (**A**) Blood Phe concentrations in male (selection n = 5; no selection n = 6) and female mice (selection n = 3; no selection n = 3) of the *Pah^enu2/enu2^*strain treated with hepatocyte transplantation. Arrows indicated when APAP diet treatment was commenced and discontinued. Dashed line indicates therapeutic threshold of 360 µM. (**B**) Coat color of female mice: wild-type C57BL/6 (WT), *Pah^enu2/enu2^* treated with cell transplant and APAP selection, *Pah^enu2/enu2^*treated with cell transplant without APAP selection, and untreated *Pah^enu2/enu2^*. (**C**) Percent Cypor-deficient hepatocytes estimated based on indel analysis from whole liver homogenate gDNA from Phe-corrected, APAP selected *Pah^enu2/enu2^*mice (n = 7) and unselected controls (n = 9). Filled circles indicate males, unfilled circles indicate females. (**D**) tdTomato fluorescence (red), Cypor immunofluorescence (green), and nuclear Hoechst (blue) in liver from an APAP-selected, Phe-corrected *Pah^enu2/enu2^* mouse (top), and an unselected control (bottom). Arrowhead indicates an individual donor hepatocyte in the unselected liver. Scale bars = 200 µM. (**E**) Pah enzyme activity in liver homogenate from treated animals as a percentage of wild-type Pah activity. Filled circles indicate males, unfilled circles indicate females. For all statistical analysis, \**P* < 0.05, \*\**P* < 0.01, and \*\*\**P* ≤ 0.001. Data are reported as means ± SD. See also Figure S2.

Analysis of indels at the Cypor sgRNA target locus in liver gDNA from corrected mice indicated an average of 13.6% ± 6.4% Cypor deficient hepatocytes, compared to 1.6% ± 1.0% in mice that received transplantation without APAP selection (p < 0.0001). Complete correction of blood Phe levels was seen in animals with a replacement index of as low as 8% (Figure 1C). Clonal expansion of Cypor-negative cells was observed in livers of APAP-treated animals by immunofluorescence (IF) staining (Figure 1D). As expected, Cypor-negative cells were positive for membranous tdTomato, indicating that they originated from the donor mouse. Pah enzyme activity assessment showed 15.4% ± 6.7% of wildtype Pah activity in corrected *Pah^enu2/enu2^* animals compared to 0% in unselected controls (p < 0.0001) (Figure 1E).

Mice of a newly described Pah-null strain(20), *Pah^Δexon1/Δexon1^*, were also treated by hepatocyte transplantation and APAP selection with the same protocol. *Pah^Δexon1/Δexon1^*animals showed similar correction, with average post-treatment blood Phe levels of 156 µM following 50-75 days of dietary treatment (Figure S3A). Coat color darkening (Figure S3B) was also observed in these mice. Indel analysis on livers of treated animals showed an average of 15.0% ± 2.8% Cypor-deficient hepatocytes, compared to 1.2% ± 1.0% in unselected animals (p = 0.0037) (Figure S3C), and clonal expansion of Cypor-negative cells was evident by IF staining (Figure S3D). IF staining for Pah in the livers of corrected *Pah^Δexon1/Δexon1^* animals revealed that Pah expression co-localized with tdTomato-positive transplanted hepatocytes (Figure S3E). IF staining for Pah is not possible in the *Pah^enu2/enu2^*model due to presence of mutant protein. Pah enzyme activity in corrected *Pah^Δexon1/Δexon1^* mice showed an average of 17.5% ± 1.2% of wild-type Pah activity compared to an average of 0.06% in unselected mice (p = 0.0001) (Figure S3F).

### Safety Outcomes

A potential safety concern for the APAP selection system is whether partial Cypor deficiency in the liver will lead to a metabolic phenotype. A mouse model of complete hepatic Cypor deficiency has been described(21, 22). Circulating levels of cholesterol and triglycerides are decreased in this mouse, and accumulation of lipids in hepatocytes corresponds to a significant increase in liver to body weight ratio. Here, corrected *Pah^enu2/enu2^* mice with only ∼14% Cypor-null hepatocytes (partial Cypor deficiency) showed no significant difference in blood levels of cholesterol or triglycerides compared to unselected controls (Figure 2A-B). Liver function markers were also measured and showed no significant differences between selected and unselected *Pah^enu2/enu2^* mice (Figure S4). Liver weight to body weight ratio also showed no significant difference between selected and unselected *Pah^enu2/enu2^* animals (Figure 2C). H&E staining of the livers of APAP selected mice revealed no distinct differences between APAP-treated and unselected animals (Figure 2D). Notably, while hepatic Cypor-null mice have hepatic accumulation of lipids, hepatic lipid accumulation was not seen in any cells in selected animals with partial Cypor deficiency of either the *Pah^enu2/enu2^* or the *Pah^Δexon1/Δexon1^*strain (Figure 2D).

**Figure 2:**
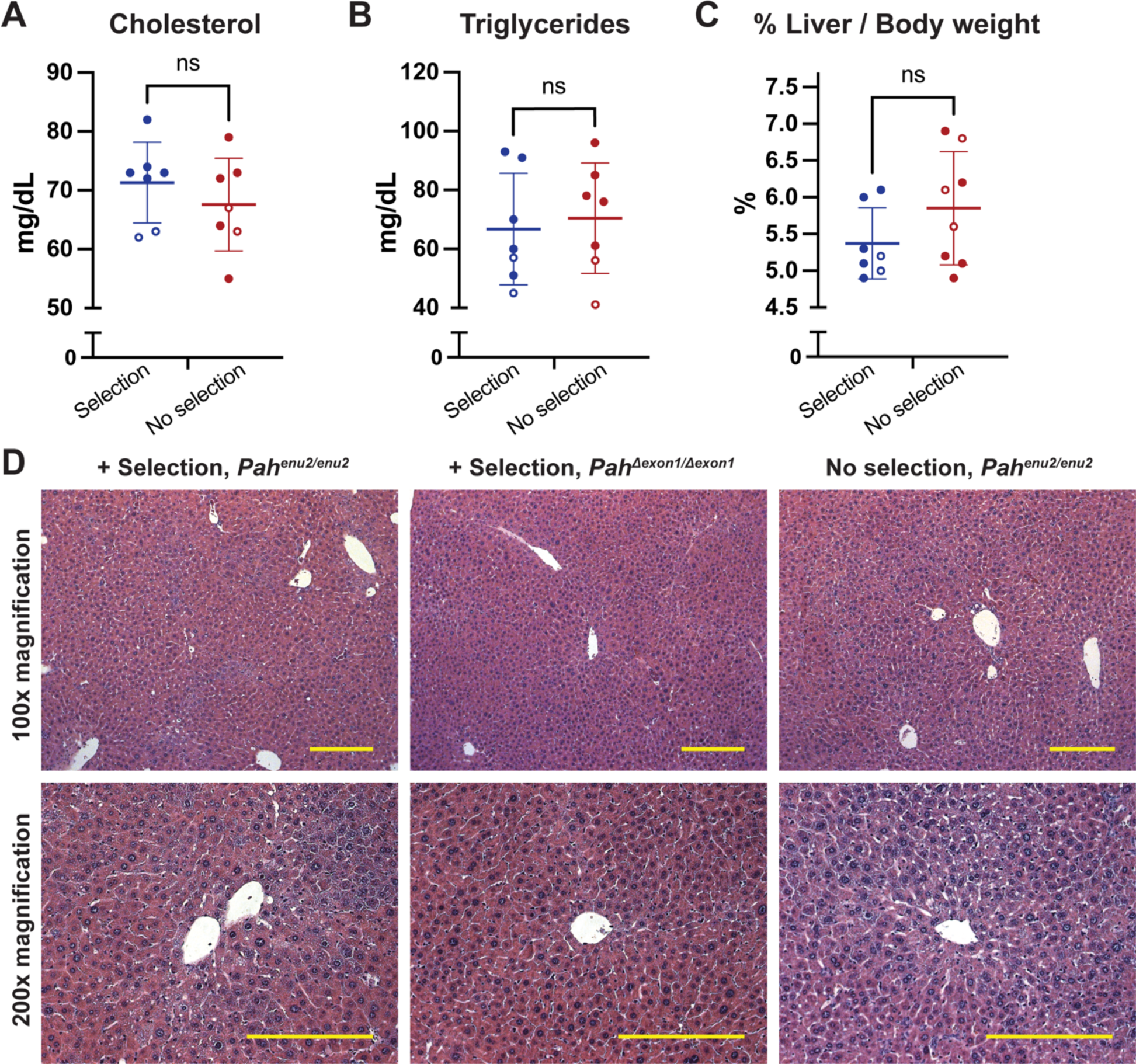
Safety assessments in corrected *Pah^enu2/enu2^* mice. (**A-B**) Blood chemistry analysis upon terminal harvest in corrected vs. unselected *Pah^enu2/enu2^*animals: (**A**) Cholesterol (**B**) Triglycerides. (**C**) Liver weight as percent of body weight in corrected vs. unselected *Pah^enu2/enu2^*animals. Filled circles represent males, unfilled circles represent females. Data are reported as means ± SD. (**D**) Representative H&E staining in a corrected *Pah^enu2/enu2^* animal, a corrected *Pah ^Δexon1/Δexon1^* animal, and an unselected control *Pah^enu2/enu2^*animal. Top: 100x magnification; bottom: 200x magnification. Scale bar = 200 µM. See also Figure S4.

### Drug metabolism

Cypor deficiency in APAP selected animals is limited to cells in zone 3 of the hepatic lobule, constituting approximately one-third to one-half of all hepatocytes, because these are the only hepatocytes that express those Cyp enzymes (Cyp2E1, Cyp1A2, and Cyp3A4) necessary for the conversion of APAP to NAPQI(8). As zone 3 Cyp activity is important for metabolism of some xenobiotic substrates(23), it is desirable to achieve the therapeutic threshold while still retaining a population of hepatocytes in zone 3 with Cypor activity. We previously performed proof of concept experiments showing that up to 40% of the liver can be made Cypor-deficient with extensive APAP selection(7). By comparison, corrected PKU animals with ∼14% Cypor deficient hepatocytes are expected to retain more than half of residual zone 3 hepatocytes with Cypor activity. Immunofluorescent staining for Cyp2E1, one of the zone 3-expressed Cyps involved in APAP metabolism(8), confirmed that the majority of Cyp2E1-expressing cells did not overlap with the tdTomato-positive transplanted cells, indicating that most zone 3 hepatocytes retain Cypor activity (Figure 3A).

**Figure 3:**
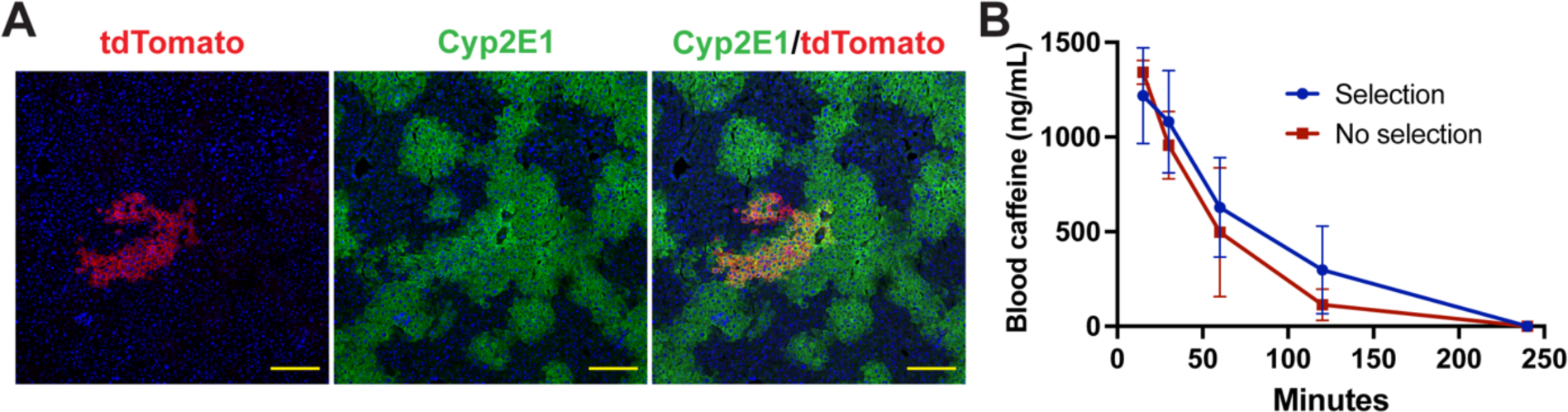
Drug metabolism in corrected *Pah^enu2/enu2^* mice. (**A**) tdTomato fluorescence (red), immunofluorescent staining for Cyp2E1 (green), and nuclear Hoechst (blue) in a corrected *Pah^enu2/enu2^* animal. (**B**) Blood caffeine concentration over time in corrected male *Pah^enu2/enu2^* mice (n = 4) and unselected control animals (n = 3). Caffeine was administered at a dose of 1 mg/kg at timepoint 0. Data are reported as means ± SD.

As a functional assay of Cypor activity, the metabolism of a Cyp-metabolized drug was assessed in selected mice. Caffeine is a well-characterized probe drug(24) for the activity of Cyp1A2, a zone 3-specific Cyp enzyme that also plays a role in the metabolism of APAP(25). It is estimated that Cyp1A2 is responsible for >95% of caffeine metabolism(26), and mice completely lacking Cypor expression in the liver have been reported to have highly deficient caffeine elimination (27, 28). As an assessment of zone 3 Cyp activity, caffeine was administered to a subset of corrected *Pah^enu2/enu2^* male mice (average 17% Cypor-deficient hepatocytes) and unselected *Pah^enu2/enu2^* male control animals. Following an intraperitoneal administration of 1 mg/kg caffeine, blood was collected at 15 minutes, 30 minutes, 1 hour, 2 hours, and 4 hours for assessment of caffeine levels. No significant difference was seen in AUC between selected and unselected animals (p = 0.34) (Figure 3B), indicating retention of adequate zone 3 Cypor activity despite partial genetic deficiency.

## Discussion

Here, we report complete, long-lasting correction of hyperphenylalaninemia in two different mouse models of PKU via selective expansion of transplanted donor hepatocytes. Hepatocyte transplantation carries several potential advantages as compared to viral gene therapy methods, which are already widely used clinically. First and foremost, wildtype hepatocytes represent a single “drug” for many different genetically distinct hepatic disorders. The donor cell expansion strategy presented here is likely to be applicable to any genetic disorder that can be corrected by expression of the wildtype gene in approximately 40% of hepatocytes. In contrast, viral gene therapy requires customization of genetic cargo to for each specific disorder, and each viral vector must go through clinical trials and be approved as a novel therapeutic. The versatility of this single therapeutic strategy may be of particular benefit for very rare disorders for which development of a novel drug for very few patients may not be economically feasible. In addition, concerns related to activity of the viral vectors in off-target tissue types are avoided, as are potentially life-threatening immune responses to high doses of viral vectors(29). While immune responses to viral vectors typically prevents redosing due to the formation of high levels of neutralizing antibodies to the viral capsid, repeat administration of hepatocytes is feasible and has been done successfully in a clinical setting(30). Applications of hepatocyte transplantation in humans have shown a favorable safety profile, including in even very young pediatric patients(1, 6). While episomal adeno-associated viral vector gene therapies have only transient efficacy in the pediatric liver due to episome loss with cell division(31, 32), transplanted hepatocytes are expected to divide with the growing liver and provide life-long benefit.

Despite its potential advantages, clinical application of hepatocyte transplantation has lagged behind gene therapy approaches due to three major obstacles: low cell engraftment levels, lack of donor material, and allogeneic rejection. The selection strategy presented here represents a way to overcome one of these problems, the low cell replacement index.

In our previous paper, we achieved near-complete correction of blood Phe in male *Pah^enu2/enu2^* animals and incomplete correction in female animals by expanding hepatocytes harboring a PAH-expressing lentivirus to >40% of the liver by APAP selection(7). In contrast, we were able to obtain complete correction of Phe levels here in both male and female *Pah^enu2/enu2^* mice following selective expansion of Pah-expressing transplanted hepatocytes with a replacement index as low as 8% of total hepatocytes. The difference in the percentage of Pah expressing hepatocytes needed to achieve correction is likely to be due to the known dominant negative effect of mutant Pah monomers. In our lentiviral gene therapy work(7), Pah enzyme activity was less than 4% although >30% of hepatocytes expressed Pah. In contrast, much higher Pah activity was achieved here, and the enzyme activity correlated very well with the cell replacement index (∼15%) in our cell transplantation experiments. Pah functions as a homotetramer, and the *Pah^enu2/enu2^*mouse expresses a mutant protein product that can assemble with the wildtype protein and inhibit the activity of the tetramer(33). It is currently unknown how many human *PAH* mutations may be similarly problematic for gene therapy. This dominant interference mechanism does not occur in cell therapy, because mutant and wildtype Pah reside in separate cells. The similar cell replacement threshold required to correct two different PKU mouse models highlights the mutation-agnostic nature of the hepatocyte transplantation strategy.

A concern for the safety of our APAP selection system is that the partial CYPOR deficiency could result in unintended consequences. Humans with germline CYPOR deficiency are affected by Antley-Bixler syndrome. The primary phenotype of this condition results from disordered skeletal development and steroid synthesis in extrahepatic tissues, and affected patients do not have a reported liver phenotype(34). Although decreased rates of metabolism of Cyp-metabolized drugs have been reported in affected patients(35), our experiments indicate that the limited Cypor deficiency created by APAP selection will not have a significant effect on drug metabolism. As human Cyp metabolism is already highly polymorphic(36), slight fluctuations are not likely to create significant medical risk.

Mice with a complete hepatic deficiency of Cypor(21, 22) have been shown to develop and reproduce normally. None of the physiologic changes that have been reported in these mice, including alterations to hepatic and circulating lipid levels, were seen in corrected PKU mice with partial Cypor deficiency. Of the Cyp-metabolized drugs that have been studied in this mouse model, caffeine is among the most affected(27). Here, AUC values for caffeine metabolism were not significantly altered as compared to unselected controls, demonstrating the metabolic safety of a limited number of hepatocytes being Cypor deficient.

As zone 1 and 2 hepatocytes are not susceptible to APAP-induced liver injury, completely ablating hepatic Cyp metabolism is not possible with our selection system. Furthermore, most major drug-metabolizing Cyps have intestinal as well as hepatic expression(37), representing another reservoir of Cyp activity that is not susceptible to depletion by APAP selection.

APAP selection was accomplished using gradual APAP administration via dietary consumption. This protocol is advantageous as it accomplishes selective expansion of hepatocytes while avoiding acute liver injury associated with APAP administration by injection(7). We envision that a clinical protocol would involve a similarly gradual APAP administration regimen. If ALT responses exceed an acceptable threshold, intervention with the antidote N-acetyl cystine would be available.

Taken together, the data presented here, indicate that APAP selection of transplanted hepatocytes is efficacious and safe in mice and that the technology was significant potential for clinical development.

## Materials and Methods

### Cypor knockout in vitro

Hepatocytes were isolated from 6 -12-week-old female mTmG mice on a C57BL/6 background by collagenase perfusion as previously described(10). A chemically modified synthetic single guide RNA (sequence: 5’-UCGUGGGGGUCCUGACCUAC-3’) targeting the mouse Cypor gene was obtained (Synthego). Along with spCas9 protein (Integrated DNA Technologies), sgRNA was delivered to primary hepatocytes using the CRISPRMAX reagent (Thermo Fisher Scientific) with modifications to manufacturer protocol. Briefly, the assembled ribonucleoprotein complexes were added to non-attached primary hepatocytes in suspension at 1x10^6^ cells / mL in HCM media (Lonza). Cells were incubated with rocking at 37°C for 2 hours before transplantation. Hepatocyte transplantation was conducted by injecting 5x10^5^ hepatocytes into the spleen. An aliquot of transfected cells was plated on a collagen-coated plate in HCM media and harvested at day 5 post-transfection for gDNA extraction and indel assessment.

### Animal husbandry

mTmG mice on the C57BL/6 background(16) (stock no. 007576) were obtained from the Jackson laboratory. *Pah^enu2/enu2^* mice(19) and *Pah^Δexon1/Δexon1^* mice(20) on a C57BL/6 background were obtained from the Harding Lab. All animal experiments were performed according to the guidelines for animal care at Oregon Health & Science University (OHSU). All animals were fed tap water and standard mouse chow (LabDiet PicoLab Rodent Diet 5LOD; ≥23% protein) unless otherwise stated. Animals were housed under a standard 12-hour on-and-off light cycle. The APAP diet was made by dissolving APAP (Sigma-Aldrich) in 20 ml of 100% ethanol and adding 180 g of standard 5LOD diet preheated to 55°C. The chow was stirred until pellets were saturated with ethanol and then allowed to air-dry at room temperature completely before being fed to mice. All mice were started on 1.2% (w/w) APAP diet at first and transitioned to a 1.5% diet after one week. Mice were given 2 to 4 days on regular diet if weight loss exceeded 20%. Mice were fasted for 4 hours prior to harvest and were euthanized with CO_2_. A cardiac puncture was performed immediately after death.

### Pah enzymatic activity and serum Phe

Liver Pah enzyme activity and serum Phe were determined as previously described(38).

### Blood Chemistry

Serum was obtained by terminal cardiac puncture and submitted to IDEXX Laboratories (www.idexx.com) for liver and lipid blood chemistry panels.

### TIDE analysis of CRISPR/Cas9-induced indels

Genomic DNA extraction from homogenized liver tissue was performed using the MasterPure Complete DNA and RNA Purification Kit (Lucigen) following the manufacturer’s protocol. An 800–base pair region surrounding the target site of the Cypor sgRNA was amplified (forward primer 5′-GTTTGCGGGTGTTAGCTCTTC-3′; reverse primer 5′-TTGGTGGGTAAATCACACCGT-3′) using MyTaq Red Mix (Bioline). The amplicon was purified using the PCR clean-up and gel extraction kit (Macherey-Nagel) and Sanger sequenced using the forward primer. Indels were analyzed using the TIDE software (https://tide.nki.nl)(17). The percentage of Cypor-deficient hepatocytes was estimated assuming that hepatocytes account for 60% of total liver DNA (percent indels ÷ 0.6 = percent Cypor-deficient hepatocytes).

### Immunohistochemistry

Approximately 5 mm-thick cross-sections of liver tissue were fixed in 4% paraformaldehyde (Sigma-Aldrich) at room temperature for 4 hours or at 4°C overnight. Liver slices were then passed through a sucrose gradient consisting of 10, 20, and 30% sucrose (w/v) in phosphate-buffered saline (PBS). Tissues were embedded in O.C.T. Compound, and 7 μM sections were cut using a cryostat onto Colorfrost Plus slides (Thermo Fisher Scientific). Sections were permeabilized in 0.25% Triton X-100 in PBS at room temperature for 12 min, washed in 3× 5-min in PBS, and then blocked in 10% normal donkey serum in PBS for 30 min at room temperature. Slides were incubated with primary antibody for 1 hour at room temperature or overnight at 4°C. The following primary antibodies were used: rabbit anti-Cypor (Abcam, no. 180597), rabbit anti-Cyp2E1 (Abcam, no. 28146), rabbit anti-PAH (Boster Bio, no. A00761-1). Slides were washed in 3× 5-min in PBS, incubated in secondary antibody (Alexa Fluor® 647-conjugated donkey anti-rabbit; Jackson Immunoresearch, no. 711-606-152) for 1 hour at room temperature, incubated in 1 µg/mL Hoechst 33342 (Invitrogen) in PBS for 3 minutes, and then washed 3× 5 min. Coverslips were mounted with Fluoromount-G (SouthernBiotech). Imaging was performed on a Zeiss LSM 700 confocal microscope.

### Blood Caffeine Analysis

Caffeine (Sigma-Aldrich) in saline was administered to mice intraperitoneally at a dose of 1 mg/kg body weight. Blood was taken from the saphenous vein at 15 minutes, 30 minutes, 1 hour, 2 hours, and 4 hours after injection. Serum was analyzed for caffeine by ELISA (Abcam, no. 285229) with modifications to manufacturer protocol. Caffeine administration was done on day 200-250 after the discontinuation of APAP diet.

### Statistical analysis

All statistical analysis was performed using GraphPad Prism version 10.0.0 for Mac (GraphPad Software, La Jolla, California, www.graphpad.com). A Student’s unpaired one-tailed *t* test assuming equal variance was used to analyze differences in percent Cypor-deficient hepatocytes, Pah activity, and body weight between selected and unselected mice. A Student’s unpaired two-tailed *t* test assuming equal variance was used to analyze differences in blood chemistry and liver to body weight ratio between selected and unselected mice. For analyzing differences in blood Phe levels between selected and unselected mice over a time course, a two-way repeated-measures ANOVA or mixed effects model with Bonferroni multiple comparisons was used. *P* values <0.05 were considered statistically significant. Area under the curve (AUC) values for caffeine metabolism (assuming baseline = 0) were computed using GraphPad Prism, followed by a Student’s unpaired two-tailed *t* test assuming equal variance. For all statistical analysis, \**P* < 0.05, \*\**P* < 0.01, and \*\*\**P* ≤ 0.001. All error bars indicate SD.

## Data availability statement

All data associated with this study are present in the paper or the supplemental materials.

## Author contributions

A.V.: Conceptualization, Investigation, Formal analysis, Visualization, Writing – Orginal draft. L.W.: Investigation. M.M.: Investigation, Formal analysis. C.O.H.: Supervision, Writing – Review & editing. M.G.: Conceptualization, Supervision, Writing – Review & editing. A.T.: Investigation, Supervision, Writing – Review & editing.

## Acknowledgements

We thank S. Winn and S. Dudley for animal husbandry and technical support, D. Enicks for assistance with cell transplants and animal husbandry and A. Major at Baylor College of Medicine for histology.

---

**Supplemental figure 1:**
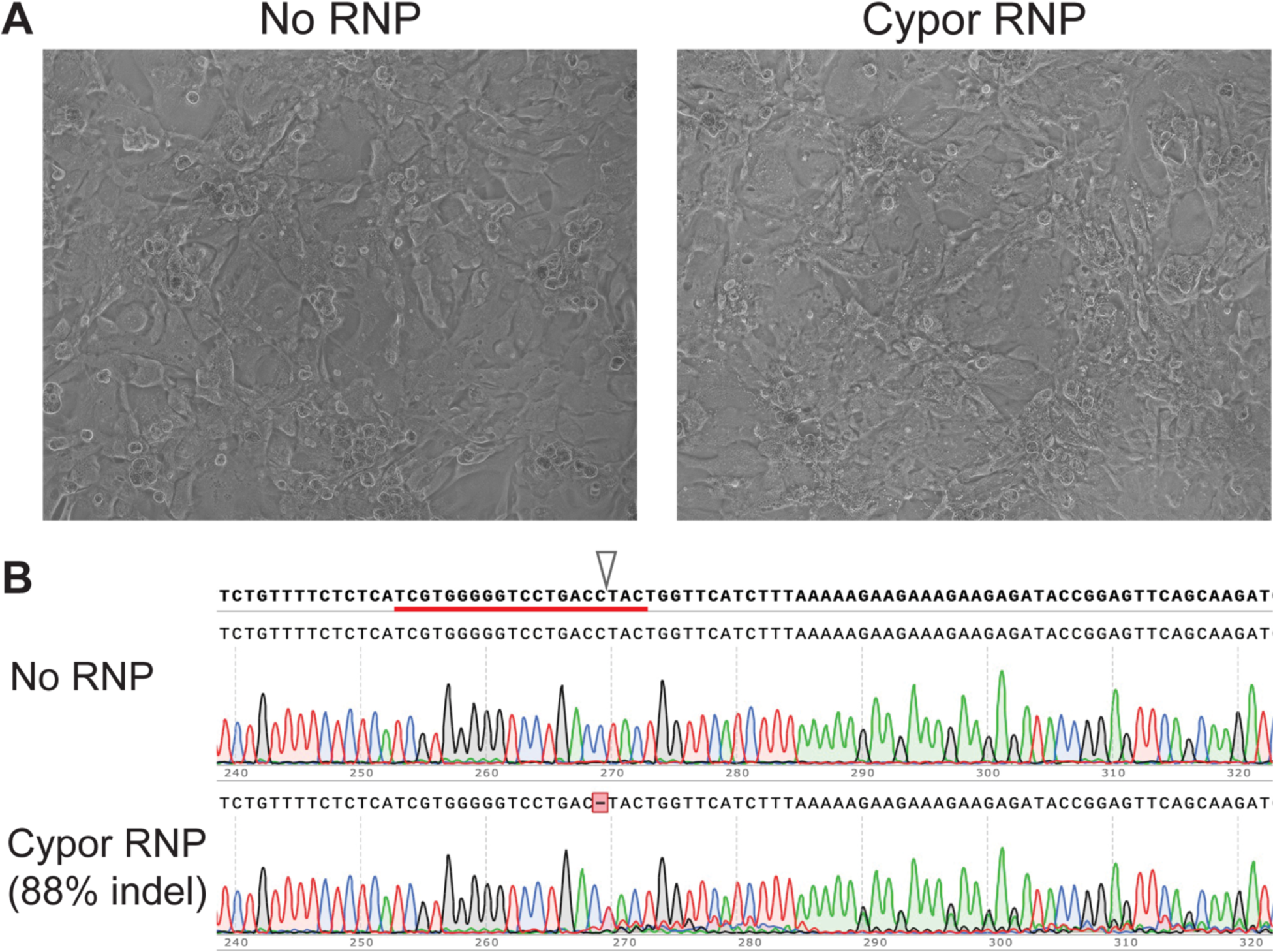
In vitro manipulation of primary mouse hepatocytes. **(A)** Plated untreated control and Cypor RNP-treated primary mouse hepatocytes on day 5 after perfusion and transfection. **(B)** Representative Sanger sequencing trace from amplicon sequencing from plated cells harvested on day 5 after transfection. gRNA sequence is underlined, and arrow indicates expected cut site. Indel percentage, as estimated from the trace using the TIDE algorithm, is indicated.

**Supplemental figure 2:**
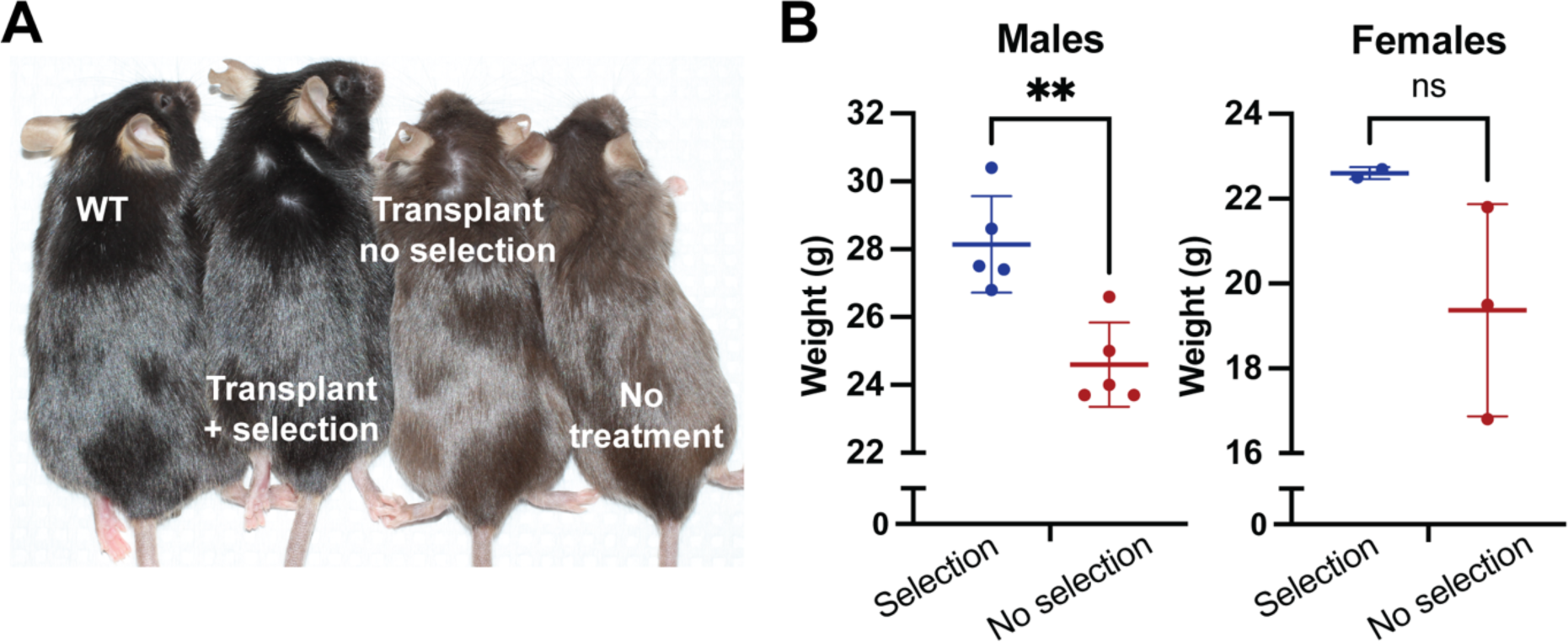
Additional efficacy data in selected *Pah^enu2/enu2^* mice. (**A**) Coat color of male mice: wild-type C57BL/6, *Pah^enu2/enu2^* treated with cell transplant and APAP selection, *Pah^enu2/enu2^* treated with cell transplant without APAP selection, and untreated *Pah^enu2/enu2^*. (**B**) Body weight at time of harvest in corrected mice and unselected controls. Error bars indicate SD.

**Supplemental figure 3:**
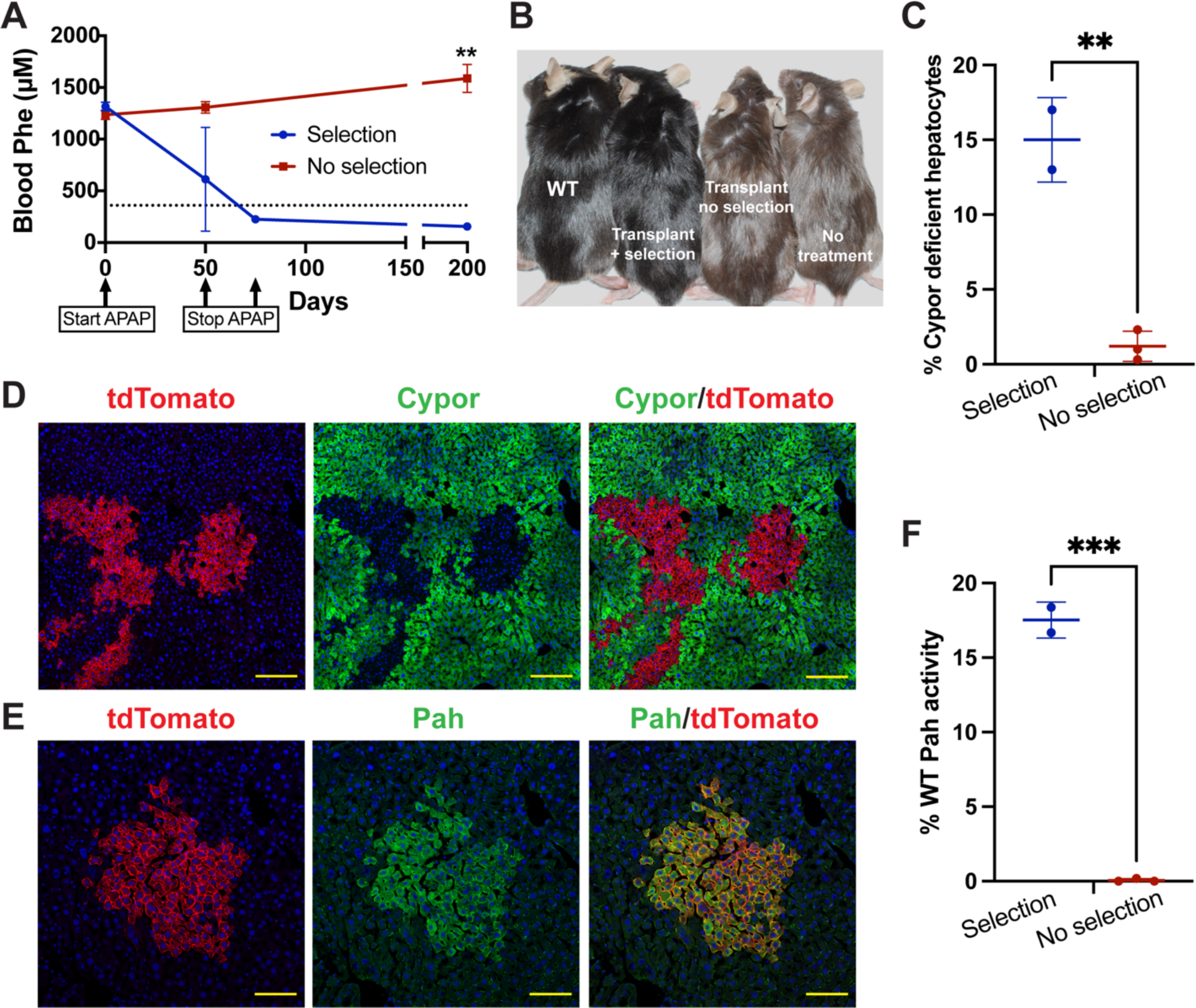
APAP selection of transplanted hepatocytes in *Pah^Δexon1/Δexon1^* mice. (**A**) Blood Phe levels in treated male mice (n = 2) and unselected controls (n = 3) of the *Pah^Δexon1/Δexon1^* strain. Dashed line indicates therapeutic threshold of 360 µM. (**B**) Coat color: wild-type C57BL/6, *Pah^Δexon1/Δexon1^* treated with cell transplant and APAP selection, *Pah^Δexon1/Δexon1^* treated with cell transplant without APAP selection, and untreated *Pah^Δexon1/Δexon1^*. (**C**) Indel analysis of corrected and unselected control livers. (**D**) tdTomato fluorescence (red), Cypor immunofluorescence (green), and nuclear Hoechst (blue) in liver from an APAP-selected, Phe-corrected *Pah^Δexon1/Δexon1^*mouse. Scale bars = 200 µM. (**E**) Pah immunofluorescence (green), tdTomato fluorescence (red), and nuclear Hoechst (blue) in liver from an APAP-selected, Phe-corrected *Pah^Δexon1/Δexon1^* mouse. Scale bars = 100 µM. (**F**) Pah activity in liver homogenate from treated *Pah^Δexon1/Δexon1^*animals as a percentage of wildtype Pah activity. All error bars indicate SD.

**Supplemental figure 4:**
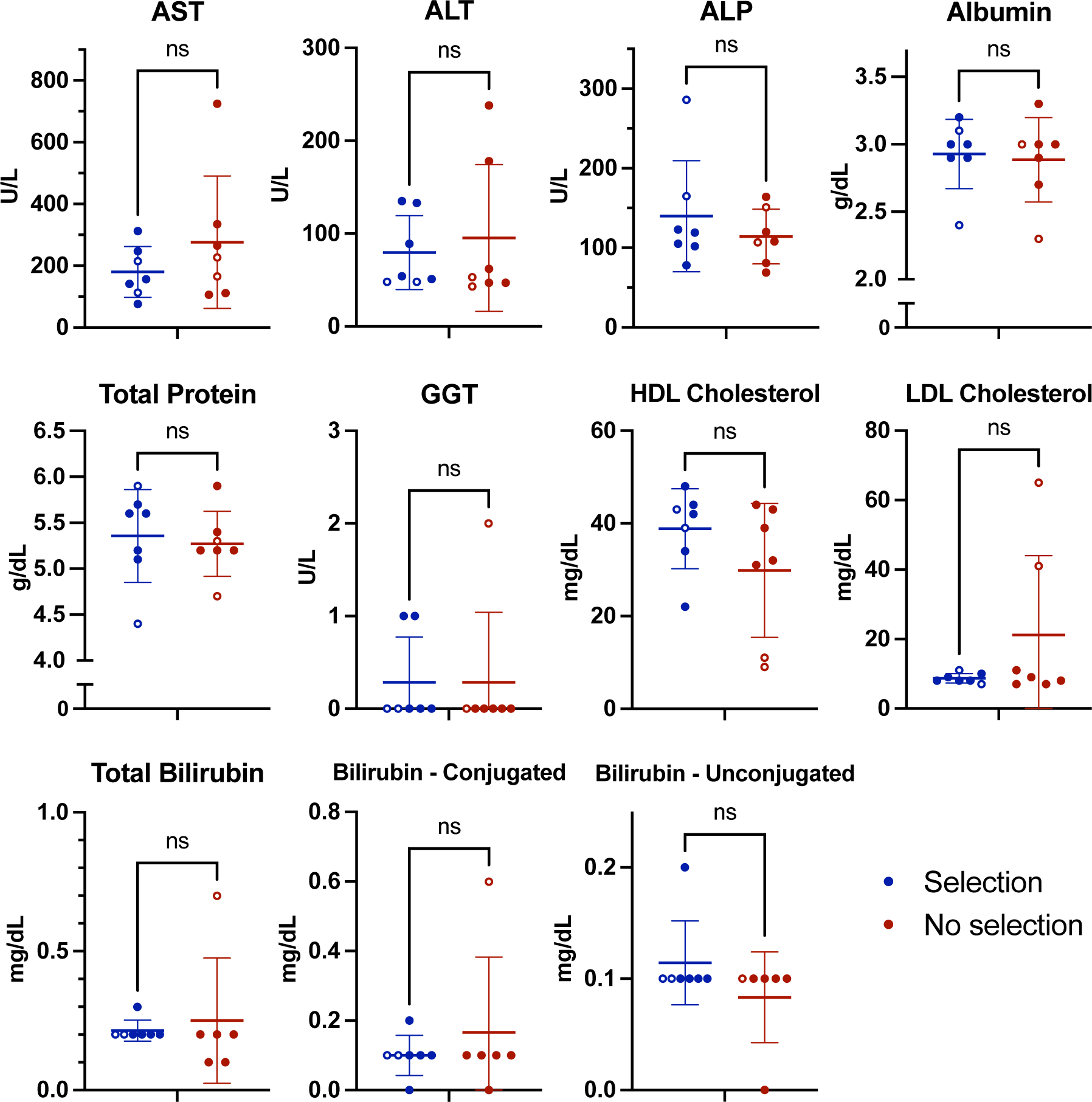
Blood Chemistry of Corrected *Pah^enu2/enu2^* mice. Liver function and lipid analysis on blood taken at terminal harvest from corrected *Pah^enu2/enu2^* animals and unselected controls. Filled circles indicate males, unfilled circles indicate females. All error bars indicate SD.

